# CellSighter – A neural network to classify cells in highly multiplexed images

**DOI:** 10.1101/2022.11.07.515441

**Authors:** Yael Amitay, Yuval Bussi, Ben Feinstein, Shai Bagon, Idan Milo, Leeat Keren

## Abstract

Multiplexed imaging enables measurement of multiple proteins *in situ*, offering an unprecedented opportunity to chart various cell types and states in tissues. However, cell classification, the task of identifying the type of individual cells, remains challenging, labor-intensive, and limiting to throughput. Here, we present CellSighter, a deep-learning based pipeline to accelerate cell classification in multiplexed images. Given a small training set of expert-labeled images, CellSighter outputs the label probabilities for all cells in new images. CellSighter achieves over 80% accuracy for major cell types across imaging platforms, which approaches inter-observer concordance. Ablation studies and simulations show that CellSighter is able to generalize its training data and learn features of protein expression levels, as well as spatial features such as subcellular expression patterns. CellSighter’s design reduces overfitting, and it can be trained with only thousands or even hundreds of labeled examples. CellSighter also outputs a prediction confidence, allowing downstream experts control over the results. Altogether, CellSighter drastically reduces hands-on time for cell classification in multiplexed images, while improving accuracy and consistency across datasets.

## Introduction

The spatial organization of tissues facilitates healthy function and its disruption contributes to disease ^1^. Recently, a suite of multiplexed imaging technologies has been developed, which enable measurement of the expression of a continuously-increasing number of proteins in tissue specimens at single-cell resolution while preserving tissue architecture ^2,3,12–14,4–11^. These technologies hold promise to fully capture the complexity of tissues and open new avenues for large-scale molecular analysis of human development, health and disease. However, while the technologies for multiplexed imaging have experienced rapid development, with datasets now spanning thousands of images, there remains a lack of analytical tools to process the massive and complex data that they generate. To date, data analysis presents a major obstacle to broad use by the scientific community, and specifically cell classification, the task of identifying different cell types in the tissue remains an inaccurate, slow and laborious process.

Analysis of multiplexed imaging has converged on a common sequence of procedures **(Fig. 1A)**. While technologies differ in implementation, from cyclic fluorescence to mass-spectrometry, they all eventually generate a stack of images, each depicting the expression of one protein in the tissue. Initial processing includes corrections for technology-specific artifacts such as autofluorescence, image registration and noise ^15–17^. Next, clean images undergo *cell segmentation*, the task of identifying individual cells in the tissue. Recently, state-of-the-art artificial intelligence (AI) algorithms, trained on large manually-curated datasets, have automated this task and are approaching human-level performance ^18–20^. Following cell segmentation, the expression of each protein is quantified in each cell to create an *expression matrix*. This table serves as input for *cell classification*, where the type and phenotype of each cell in the tissue is inferred using the combination of co-expressed proteins, in combination with prior biological knowledge. For example, a cell expressing CD45 will be classified as an immune cell. A cell that in addition expresses CD3 and CD8 is a cytotoxic T cell, and if that cell also expresses PD-1, LAG-3 and TIM-3, it is classified as an exhausted cytotoxic T cell ^21^.

**Figure 1:**
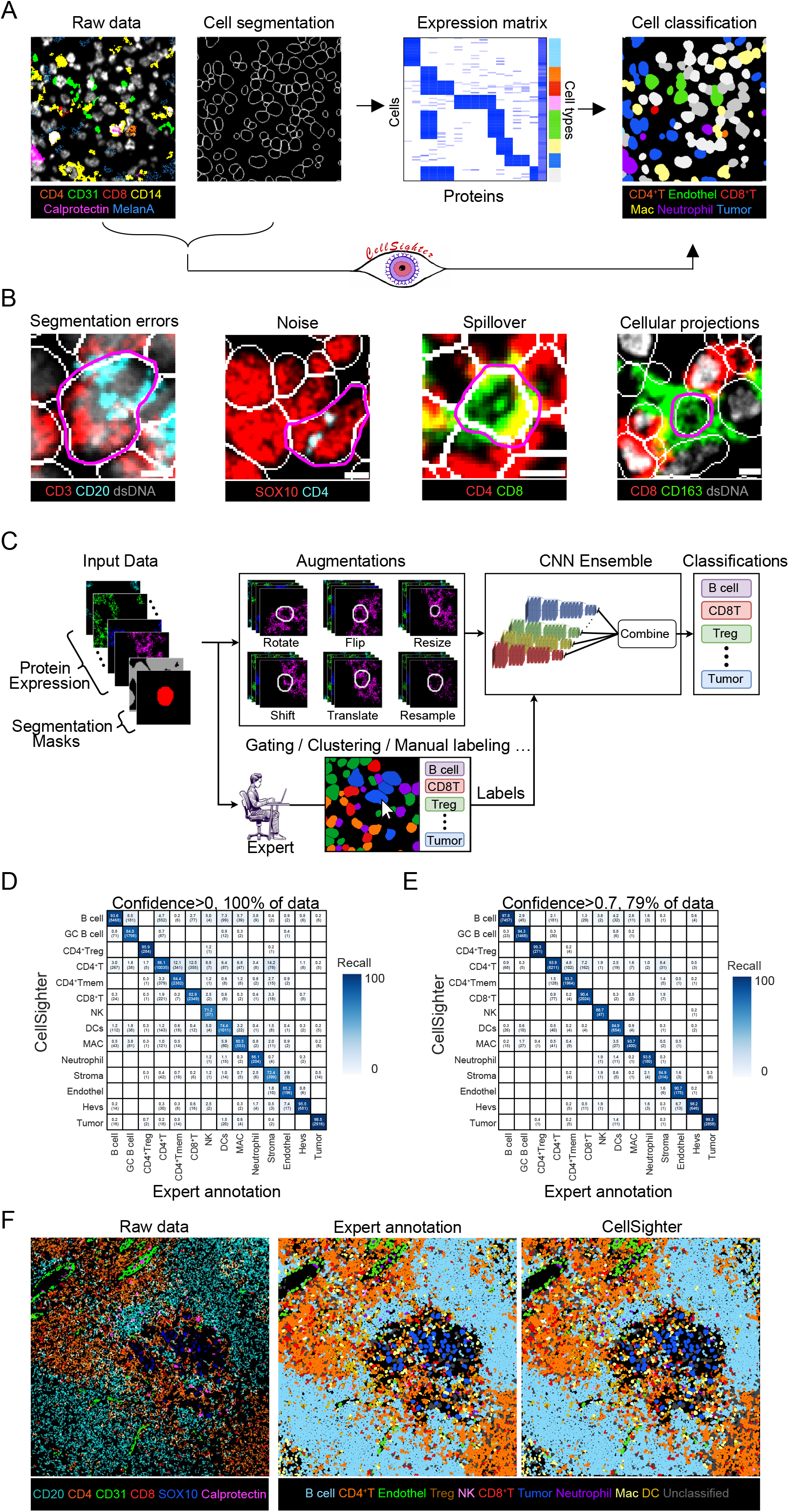
CellSighter – a convolutional neural network for cell classification. **(A)** Standard pipelines for cell classification take in multiplexed images and cell segmentation masks and generate an expression matrix, which is clustered to annotate cell types. CellSighter works directly on the images. **(B)** Imaging artifacts and biological factors contribute to making cell classification from images challenging. Segmentation errors, noise, tightly packed cells and cellular projections are easily visible in images, but hard to discern in the expression matrix. Scale bar = 5*μ*m. **(C)** CellSighter is an ensemble of convolutional neural networks (CNNs) to perform supervised classification of cells. **(D)** Comparison between labels generated by experts using clustering, gating and manual annotation (x-axis) and labels generated by CellSighter (y-axis) shows good agreement. **(E)** Same as (D) for high-confidence classifications (Confidence>0.7). **(F)** For one FOV, shown are the protein expression levels (left), expert-generated labels (middle) and CellSighter labels (right).

Cell classification methods typically involve manual gating or clustering of vectors in the expression matrix using algorithms that were developed for isolated cells, including flow cytometry, mass cytometry or single cell RNA sequencing (scRNAseq) ^5,10,17,22–28^. However, deriving cell classifications from multiplexed images has unique challenges over classifying cells in suspension, owing to both biological and technical factors. Technically, although segmentation algorithms have improved dramatically, errors in segmentation are still frequent. Fig 1B (I) shows an example of a B cell and a T cell which were erroneously segmented as one cell. In the expression matrix, this cell will appear as expressing both CD20 and CD3. Imaging artifacts including background, noise and antibody aggregates also make classification challenging ^15^. Fig 1B (II) shows an example of a tumor cell, with overlapping noise in CD4. In the expression matrix, this cell will erroneously appear to express CD4.

Biological factors also contribute to the difficulty of cell classification. In tissues, cells form densely-packed communities. For example, Fig. 1B (III) shows a cytotoxic T cell closely interacting with several T helper cells. Cells communicate with each other by interactions of ligands and receptors on their membranes and extend cellular projections to facilitate these interactions ^29^, as shown for the CD163^+^ macrophage in Fig 1B (IV). This close-network of cell bodies and projections results in *spillover*, whereby the protein signals from one cell overlap with the pixels of neighboring cells. Several works have proposed methods to deal with spillover using compensation ^3,30^, pixel analysis ^31^ or neighborhood analysis ^32^, but these suffer from signal attenuation, difficulty in scaling to large datasets, or requirements for additional data sources such as scRNAseq on the same tissue. Altogether, cell classification has hitherto remained a time-consuming and labor-intensive task, requiring sequential rounds of clustering, gating, visual inspection and manual annotation. Accordingly, the accuracy of classification is often user-dependent and may impede the quality of downstream analysis.

Here, we present CellSighter, a deep-learning based pipeline to perform cell classification in multiplexed images. Given multiplexed images, segmentation masks and a small training set of expert-labeled images, CellSighter outputs the probability of each cell to belong to each cell type. We tested CellSighter on data from different multiplexed imaging modalities and found that it achieves 80-100% accuracy for major cell types, which approaches inter-observer concordance. Ablation studies and simulations showed that CellSighter learns features of protein expression levels, but also spatial features such as subcellular expression patterns and spillover from neighboring cells. CellSighter’s design reduces overfitting and it can be easily trained on only thousands or even hundreds of labeled examples, depending on cell type. Importantly, CellSighter also outputs confidence in prediction, allowing an expert to evaluate the quality of the classifications and tailor the prediction accuracy to their needs. Finally, CellSighter can be applied across datasets, which facilitates cross-study data integration and standardization. Altogether, CellSighter drastically reduces hands-on time for cell classification in multiplexed images, while improving accuracy and consistency across datasets.

## Results

### CellSighter – a convolutional neural network for cell classification

In this work we sought to accelerate and improve cell classification from multiplexed imaging by harnessing two insights into this task. First, the effects of segmentation errors, noise, spillover and projections accumulate over the millions of cells routinely measured in multiplexed imaging datasets. As a result, the time that it takes to classify cells using manual labeling, gating and clustering is proportional to the number of cells in the dataset. It is easier and faster to classify thousands of cells than to classify millions of cells. This observation implies that a machine-learning approach that learns classifications from a subset of the data and transfers them to the rest of the dataset could largely expedite the process of cell classification. Second, while segmentation errors, noise, spillover and projections confound the protein expression values in the expression matrix, they are often distinguishable in images. We therefore reasoned that a computer-vision approach that works directly on the images as input, rather than on the expression matrix, may have better performance in the task of cell classification. Specifically, deep convolutional neural networks (CNNs) have had remarkable success in computer vision tasks and have recently gained traction in medical imaging, from radiology to electron microscopy ^18,20,33,34^.

We designed CellSighter as an ensemble of CNN models that, given raw multiplexed images, as well as the corresponding segmentation mask, returns the probability of each cell to belong to one of several classes **(Fig. 1C)**. The input for each model is a 3-dimensional tensor, consisting of cropped images of K proteins centered on the cell to be classified (**Supp. Fig. 1A, B)**. To incorporate the information of the segmentation, we added two additional images to the tensor. The first consists of a binary mask for the cell we want to classify (1’s inside the cell and 0’s outside the cell), and the second is a similar binary mask for all the other cells in the environment. To deal with large class imbalances between cell types, which in tissues can easily reach 100-fold ^24,35^ when training the network we upsampled rare cells such that the major lineages are represented in equal proportions (Methods). However, for very low-abundant classes upsampling alone can result in spurious correlations and overfitting. We therefore added standard and custom augmentations to the data, including rotations and flips, minor resizing of the segmentation mask, minor shifts of the images relative to each other, and signal averaging followed by Poisson sampling (Methods).

The final cell classification is given by integrating the results from ten separately-trained CNNs. Each model predicts the probability of each class for the cell. We then average those probabilities and take the class with the maximum probability as the final prediction and the probability value as the confidence. We found that this design provides results that are more robust and reduces grave errors of the network, including hallucinations ^36^. The confidence score gives the investigators of the data freedom to further process CellSighter’s results to decide on the level of specificity, sensitivity and coverage of cells in the dataset that are best for their specific needs. Low-confidence cells can also be used to guide and refine further labeling.

We first tested CellSighter on a dataset of human melanoma lymph node metastases, acquired by MIBI-TOF ^2^. We took sixteen 0.8×0.8mm^2^ images, encompassing 116,808 cells and generated labels for all cells using established approaches, including FlowSOM clustering ^22^, pixel clustering ^31^, gating and sequential rounds of visual inspection and manual annotation (Methods). Altogether, we distinguished fourteen cell types, including different types of tumor cells, stromal cells, vasculature and immune cells. We trained CellSighter on twelve images and tested it on four held-out images. Prediction accuracy on the test was high (85±8%) **(Fig. 1D)**, and strikingly similar to the concordance between two different human labelers (84±17%, **Supp. Fig. 1C**). It ranged from 99% on easily-distinguishable cells, such as tumor cells and high endothelial venules (HEVs) to 70% on rare, entangled or lineage-related cell types. For example, *Memory CD4 T cells* were mostly confused with lineage-related *CD4 T cells* (12%). We evaluated to what extent these confusions represent errors in CellSighter, errors in the expert annotations or ambiguous cells. To this end, we had an expert manually inspect 300 random cells that received different annotations by CellSighter and the expert, without knowing which approach provided which annotation. We found that in 30% of cases the confused cells were ambiguous and could be classified as either type, in 9% they were both wrong, in 32% manual inspection agreed with CellSighter and in 29% with the expert **(Supp. Fig. 1D)**. Overall, this suggests that discrepancies are mostly driven by ambiguous cells, and CellSighter performs comparably to current approaches. Importantly, limiting the analysis to high-confidence cells, using a cutoff of 0.7 on the probability, results in classifications for 79% of the dataset and increases the accuracy of prediction to 93±5% **(Fig. 1E)**. Altogether, visual inspection of the test images confirmed that CellSighter indeed recapitulated both the predictions of individual cells and of the tissue organization at large **(Fig. 1F)**.

### CellSighter learns protein coexpression patterns

We explored which features drive CellSighter’s predictions. First, we checked whether a CNN running on a tensor of protein images was able to learn protein expression per cell type, similar to what an expert does when working on the expression matrix. To this end, we correlated between the cellular protein expression levels and CellSighter’s confidence in prediction. For example, the confidence in predicting Neutrophil was highly correlated with the cellular expression levels of Calprotectin (R = 0.76, P < 10^−20^, **Fig. 2A**). Performing this analysis for all proteins across all cell types revealed expected associations between cell types and their respective proteins **(Fig. 2B)**. For example, the confidence of B cell classification was mostly positively correlated with the expression of CD20 (R = 0.68, P= P < 10^−20^) and to a lesser extent with CD45RA (R = 0.5, P < 10^−20^) and CD45 (R = 0.28, P= P < 10^−20^). We also found that CellSighter is aware of the problems of spillover and multi-class classification as B cell classification was also mildly negatively correlated with the expression of CD3 (R = −0.25, P < 10^−20^), CD4 (R = −0.23, P < 10^−20^) and CD8 (R = −0.19, P < 10^−20^). Indeed, a scatter plot of cellular CD20 expression (a hallmark protein for B cells) versus cellular CD4 expression (a hallmark protein for T helper cells) revealed that CellSighter was confident in its classifications for cells that had high expression of one of these proteins (mean confidence = 80%), but had lower confidence in the classification of cells that expressed both proteins (mean confidence = 48%, **Fig. 2C)**.

**Figure 2:**
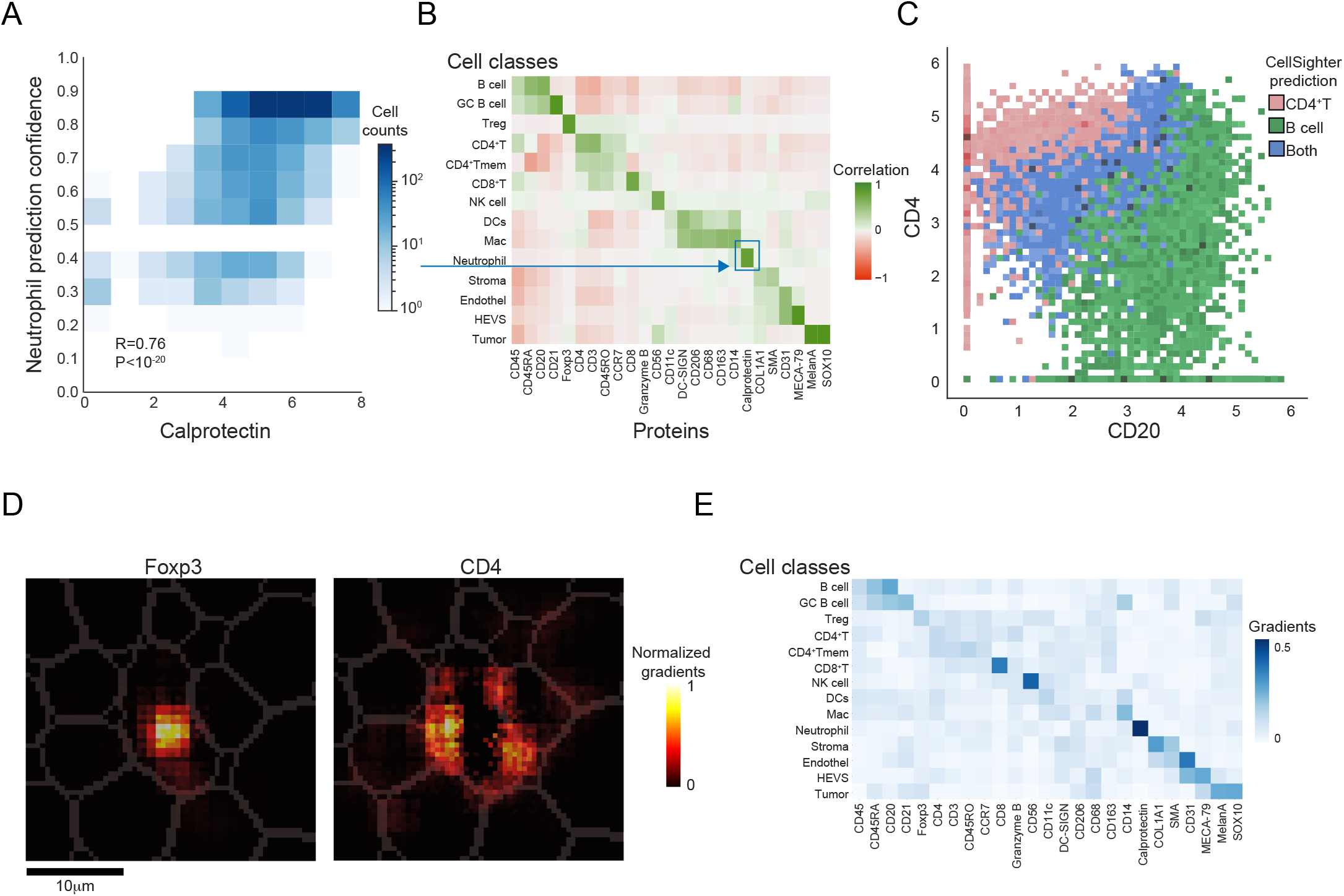
CellSighter learns protein expression patterns. **(A)** For all cells predicted as Neutrophils, shown is a 2D histogram depicting the correlation between the cellular expression levels of Calprotectin (x-axis) and CellSighter’s confidence in predicting Neutrophil (y-axis). **(B)** For each protein (x-axis) in each cell type (y-axis), shown is the correlation between the cellular expression levels of this protein and the confidence in prediction of this cell type. High positive correlations are observed between lineage proteins and their cognate cell types. **(C)** Scatter plot of cellular expression levels of CD20 (x-axis) vs. CD4 (y-axis). Cells that express mostly CD20 or CD4 were generally classified by CellSighter as B cells (green) or T helper cells (pink), respectively. Cells that express both proteins had intermediate confidence (>30%) in CellSighter’s predictions for both populations (blue). **(D)** Shown are the normalized gradients of the CNN for a single cell classified by CellSighter as a Treg, in the images of FoxP3 (left) and CD4 (right). **(E)** Shown is the average positive gradients for each protein (x-axis) in each class (y-axis) normalized across the proteins, calculated from a single CNN on 4,961 cells.

To further probe CellSighter’s classification process, we examined the gradients of the network using guided back propagation ^37^. We found that the gradients were concentrated in the center cell **(Fig. 2D)** and that they match the expected protein expression patterns. For example, in cells classified as tumor cells, the strongest gradients were traced back to the images of MelanA and SOX10, whereas for HEVs it was in the images of CD31 and MECA-79 **(Fig. 2E)**. Altogether, these analyses suggest that protein expression levels are a major determinant of CellSighter’s classification process, similar to gating and clustering.

### CellSighter learns spatial expression features

Next, we examined whether CellSighter was able to leverage the fact that it works directly on the images and learn spatial features to aid in classification. Since CellSighter was trained on data resulting from clustering the expression matrix, which suffers from spillover, we wondered whether it could generalize to learn spatial expression patterns. To do this we employed three complimentary approaches: examining the gradients, contrasting CellSighter with a machine learning model that works on the expression matrix rather than on the images, analyzing performance on simulated data, and examining the network’s gradients.

First, we evaluated whether CellSighter was more robust to spillover compared with using machine learning approaches that work on the expression matrix. To this end, we used the same set of twelve labeled images to train a gradient boosting classifier (XGBoost) that works on the expression matrix ^38^. We then used XGBoost to predict the labels for the test images and got high prediction accuracies, globally comparable to CellSighter **(Supp. Fig. 2A)**. However, visual inspection of the images suggested that CellSighter was more robust to spillover **(Fig. 3A)**. For example, inspection of Calprotectin showed that for each patch of signal CellSighter tended to classify less cells overlapping with that patch as Neutrophils **(Fig. 3A, B)**. Moreover, cells that were classified as neutrophils by CellSighter were mostly the cells that had high overlap with the signal **(>20%, Fig. 3C)**. To verify that this observation was causal, we performed a simulation where we took a patch of Calprotectin signal and moved it to vary its overall overlap with the cell **(Supp Fig. 2B)**. We found that for low overlap (<20%) CellSighter was 9% less likely to classify cells as Neutrophils compared to the XGBoost, whereas for higher degrees of overlap (>30%) this trend flipped **(Supp Fig. 2C)**.

**Figure 3:**
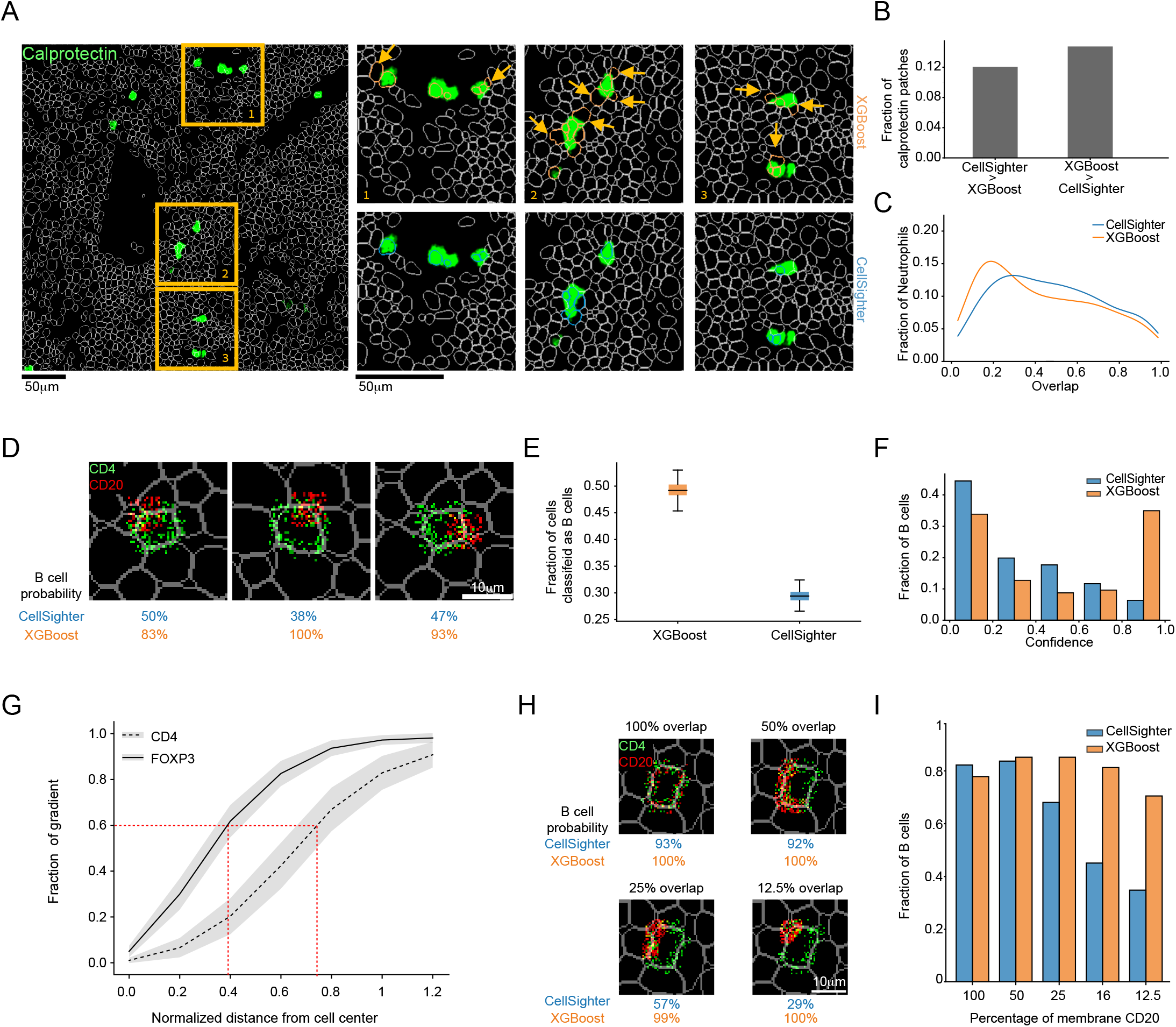
CellSighter learns spatial expression features. **(A)** Left: Calprotectin signal (green) and cell segmentation boundaries (white) in one field of view. Right: Zoom-in on boxes 1-3. The rows show cells classified as Neutrophils by either XGBoost (top row, yellow) or CellSighter (bottom row, blue). Yellow arrows point to cells that were classified as neutrophils by XGBoost, but not by CellSighter. **(B)** The number of cells classified as Neutrophils by XGBoost or CellSighter was quantified for all patches of Calprotectin signal. Bar graphs show the proportion of calprotectin signal patches (y-axis) in which the number of Neutrophils predicted was larger in either CellSighter (left) or XGBoost (right) respectively. **(C)** For all cells classified as Neutrophils in the test images, shown is the proportion of cells (y-axis) according to the level of intersection of their Calprotectin signal with the cell segmentation (x-axis). CellSighter (blue) classifies less cells with low intersection of calprotectin than XGBoost (orange). **(D)** Example images from simulations in which CD4 (green) is simulated as a membranous signal and CD20 (red) as a patch, partially overlapping with the cell membrane. **(E)** Results of simulations of 549 cells as shown in (D) randomly repeated 100 times. Shown is the fraction of simulated cells classified as B cells (y-axis) by either XGBoost (left) or CellSighter (right). **(F)** Results of simulations of 549 cells as shown in (D). Shown is the fraction of simulated cells (y-axis) as a function of the models’ confidence in B cell classification (x-axis). CellSighter (blue) has overall lower confidence in erroneous B cell classifications relative to XGBoost (orange). **(G)** Shown is the normalized sum of gradients (y-axis) as a function of the normalized integrated distance from the cell center (x-axis) for 1,353 cells classified as T helper and 42 cells classified as T regulatory cells. Gradients for the nuclear protein FoxP3 (solid line) reach 60% of their maximum at a distance of approximately 40% of the cell, whereas gradients for the membranous protein CD4 (dashed line) reach 60% of their maximum at a distance of approximately 75% of the cell. **(H)** Example images from simulations in which CD4 (green) is simulated as a membranous signal and CD20 (red) as a membranous signal with differential overlap of the membrane, ranging from 100% to 12.5%. **(I)** Results of simulations of 200 cells as shown in (H). Shown is the fraction of simulated cells classified as B cells (y-axis) as a function of the percent of the membrane that had CD20 signal (x-axis). Despite maintaining similar signal levels across the simulation, CellSighter (blue) was much less likely than XGBoost (orange) to classify cells as B cells when the overlapping segment of the membrane was small.

To further examine CellSighter’s ability to learn spatial expression patterns, we performed a direct simulation of spillover that generates contradicting expression patterns. Here, we took crops centered on T helper cells and removed their cognate CD4 and CD20 signals, to avoid any confounding factors incurred by the original signals. We then reintroduced CD4 as a membranous signal, and a patch of strong, partially-overlapping CD20 signal, to simulate spillover **(Fig. 3D)**. To verify that our simulation is relevant to real-world data, the expression levels for both CD4 and CD20 were compatible with the distribution of observed values in the dataset **(Supp. Fig. 2D)**. We then ran both CellSighter on the images and XGBoost on the expression matrix of the resulting cells. Not surprisingly, if we only added the CD4 signal, both models classified 70-80% of cells as CD4 T cells **(Supp. Fig. 2E)**. However, when introducing the CD20 signal, CellSighter classified 30% of the cells as B cells, whereas XGBoost classified 47% as B cells **(Fig. 3E)**. Moreover, CellSighter had overall lower confidence in these classifications than XGBoost. For example, CellSighter had high confidence (>0.9) that 1.5% of these simulated cells are B cells, compared to 30% for XGBoost **(Fig. 3F)**. Altogether, we conclude that both in real data and in simulations, running a CNN on images is more robust to spillover.

Next, we evaluated whether CellSighter was able to learn the sub-cellular expression patterns of different proteins. Visual inspection of the gradient maps for several cells suggested that the gradients of nuclear proteins were concentrated in the center of the cell, whereas the gradients for membrane proteins followed the segmentation borders **(Fig. 2D)**. We therefore used guided back propagation to examine the gradients of the network at varying radii from the cell center. We found that for nuclear proteins, such as FoxP3 and SOX10, CellSighter turns its attention closer to the center of the cell whereas for membrane proteins this distance increases. For example, for FoxP3, 60% of the gradient is achieved at a distance of 40% from the center of the cell, whereas for CD4 it is at 75% **(Fig. 3G, Supp. Fig. 2F)**.

To examine whether this relationship was causal we performed additional simulations. Again, we took patches centered on T helper cells and removed their cognate CD4 and CD20 signals, to avoid any confounding factors incurred by the original signals, and then reintroduced CD4 as a membranous signal. However, this time we also introduced CD20 as a membranous signal on the same cell. In different simulations we varied the percent of membrane that was covered by CD20, ranging from 100% to 12.5% **(Fig. 3H)**. We also varied the overall signal in a complimentary manner, such that the average CD20 per cell was maintained at a relatively constant level **(Supp. Fig. 2G)**. Here too we made sure that the overall signal was drawn from the real CD20 expression distribution (Methods). We found that when the CD20 signal surrounded 100% of the membrane, both models were equally likely to classify the cell as a B cell, resulting in 84% of the cells classified as B cells using CellSighter and 80% using XGBoost. However, as the fraction of overlap with the membrane was reduced, XGBoost continued to classify a similar percentage of cells as B cells, whereas CellSighter was less likely to classify the cells as B cells. For example, at 12.5% overlap, XGBoost classified 72% of the cells as B cells, whereas CellSighter dropped to 35% **(Fig. 3I)**. This suggests that CellSighter learned the characteristic expression pattern of membranous signals.

Overall, we found that CellSighter learns both protein expression levels and spatial expression patterns and integrates both when classifying cells. We note that CellSighter was able to learn these spatial features even though it was trained on imperfectly-labeled data, where annotations were mostly generated using gating and clustering on the expression matrix. This is important because most labs who perform multiplexed imaging can relatively easily generate such imperfect annotations for a subset of the data, whereas generating high-quality manually-curated annotations is difficult and time-consuming. The fact that CellSighter learns spatial and sub-cellular expression features suggests that it is able to generalize beyond just learning the clustering.

### CellSighter features contribute to performance

Next, we evaluated how different features of CellSighter affect the performance of the predictions. First, we examined what benefits, if any, are incurred by using an ensemble of models. To this end, we correlated between the cellular protein expression levels and the confidence in prediction using either a single CNN or an ensemble. We found improved correlations using the ensemble. For example, using a single CNN 2.68% of the cells that were classified as Neutrophils had low expression of Calprotectin (<2), yet they were classified as Neutrophils with high confidence (>50%). Using the ensemble, this number drops 5-fold to 0.47%, and cells that are classified as Neutrophils either have high expression of Calprotectin or are classified with low confidence **(Fig. 4A)**. Moreover, we compared the confidence in prediction for all the cells for which our prediction agreed with the expert labeling to the confidence in prediction for all the cells for which our prediction disagreed with the expert labeling **(Fig. 4B)**. We found that using a single model 20% of the cells that were wrongly classified had high confidence (>0.9). However, using an ensemble, this number dropped to 5%, and the overall confidence for wrong classifications was significantly lower **(Fig. 4B)**. Neural networks can be difficult to train and can learn spurious correlations. Using an ensemble of models provides results that are more robust, with improved correlation between the network’s confidence in prediction and its accuracy.

**Figure 4:**
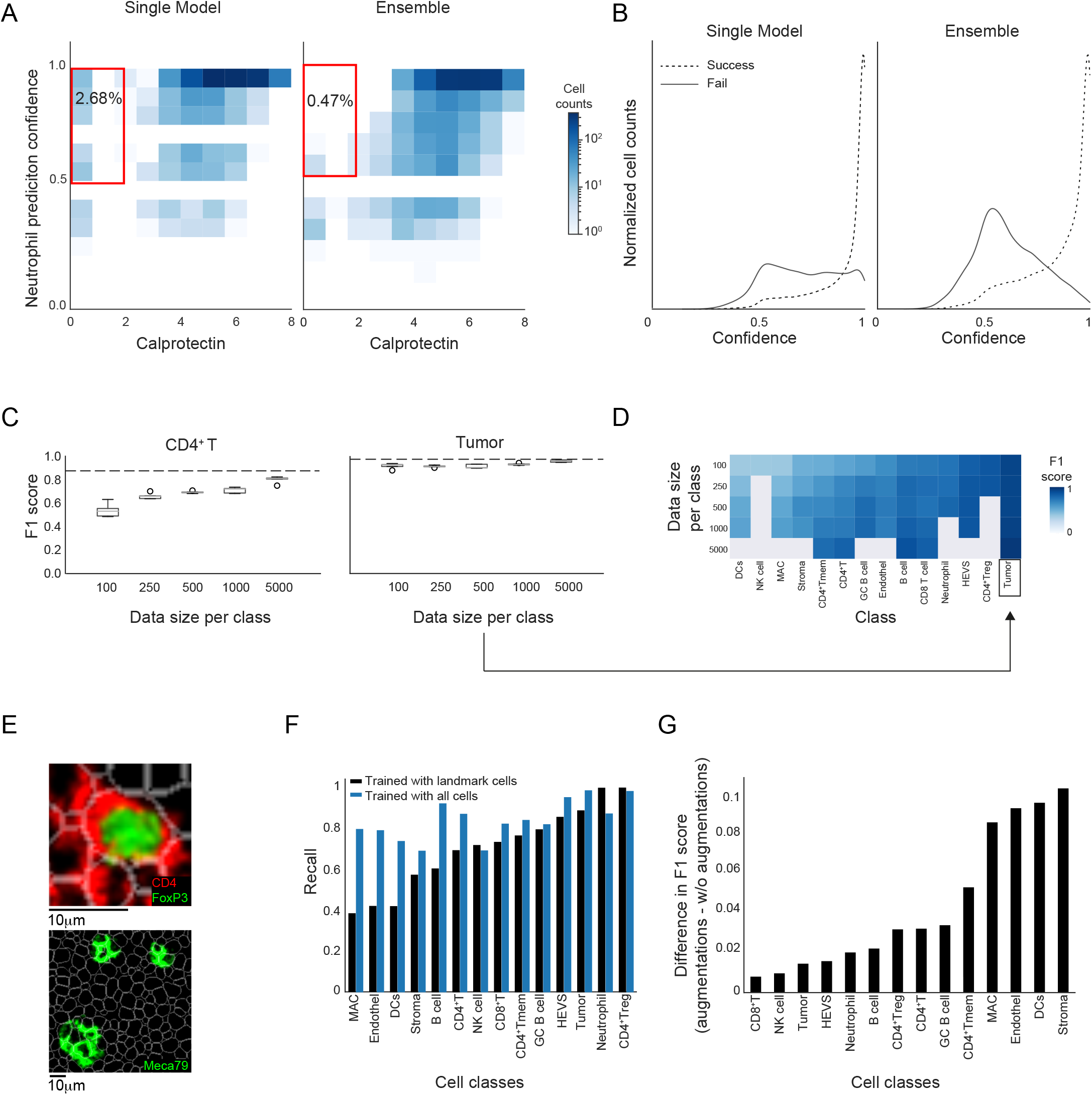
Analysis of the contribution of different CellSighter features to performance. **(A)** For all cells predicted as Neutrophils, shown is a 2D histogram depicting the correlation between the cellular expression levels of Calprotectin (x-axis) and CellSighter’s confidence in predicting Neutrophil (y-axis). Using an ensemble of models (right) reduces the fraction of cells with low Calprotectin expression that are confidently classified as neutrophils (red square) compared to a single model (left). **(B)** Shown is the fraction of cells (y-axis) for varying confidence levels (x-axis), for cells in which CellSighter predictions agree or disagree with expert labeling (solid and dashed lines, respectively). Using an ensemble (left) reduces the fraction of cells where CellSighter predicts with high confidence a class that differs from expert annotations relative to a single model (right). **(C)** For two classes, CD4 T cells (left) and Tumor (right), shown is the classification accuracy (F1 score, y-axis) as a function of the number of cells from that class used for training (x-axis). Increasing the number of cells in the training improves accuracy for CD4 T cells, but less for tumor cells. Boxplots show the results from five independent experiments with randomly sampled cells. Dashed lines indicate accuracies obtained by training on the entire training set. **(D)** Shown is the prediction accuracy (blue) for each class (x-axis) when varying the number of cells from that class in the training (y-axis). Gray colors indicate experiments that could not be performed since the dataset did not contain enough cells. **(E)** Examples of cell classes that are easily classified with a small training set, either because they rely on unique nuclear signals (Treg, top) or they organize in highly-defined spatial structures (HEVs, bottom). **(F)** Shown are classification accuracies (y-axis) for different cell classes (x-axis) using a model that was trained on landmark cells (black) or comprehensive annotations (blue). (**G)** Shown are the differences in accuracy of classification (y-axis) for different cell classes (x-axis) between a model that was trained with augmentations and a model that was trained without augmentations. The mean F1 score of epochs 35,40 and 45 is shown for each class.

In addition, we evaluated how much data is needed to train CellSighter. To this end, we retrained CellSighter on training sets that varied in size, where we randomly sampled from each class either 100, 250, 500, 1000 or 5000 cells and evaluated the resulting accuracy in prediction. To obtain confidence intervals, we repeated this process 5 times, each time training a separate model **(Fig. 4C, D)**. We found that the number of cells needed to plateau the prediction accuracy was highly variable between classes. For example, for CD4 T cells we observed a continuous improvement in F1 score from 50% to 80% when increasing the number of cells. In contrast, for tumor cells we found that increasing the training set resulted in only a modest increase in F1 score from 91% to 96% **(Fig. 4C)**. These results indicate that some cell types are easier to learn than others. This can result from these classes having better defining markers, such as nuclear proteins that are less prone to spillover. Another factor contributing to making a class more easily classifiable could be spatial organization patterns where cell types of the same class cluster together, such as in the case of HEVs **(Fig 4E)**. Overall, these results suggest that labeling efforts can be prioritized in an iterative process. A user can label a few images and train CellSighter using either all cells or a subset to identify classes that would benefit most from additional training data. The CellSighter repository supplies functions to facilitate such analyses.

Encouraged by our results that showed that CellSighter could be trained on only thousands or even hundreds of cells, we checked whether it was possible to train the network on easy, well-defined cells, that don’t suffer from segmentation errors, noise and spillover. If possible, this workflow would be highly advantageous as it would drastically reduce the time invested in the initial labeling. To this end, we identified in the dataset landmark cells that can be quickly defined using conservative gating (Methods). We then retrained CellSighter only on landmark cells from the 12 images in the training set. As expected, evaluating this model only on landmark cells in the test set achieves excellent results (97±4%, **Supp. Fig. 3A)**. Next, we tested how this model performs on the entire test set. We found that using only landmark cells for training results in good classifications for well-defined classes such as Tregs, Tumor cells and HEVs. However, overall, it achieved poorer classifications relative to using an unbiased representation of the dataset (70±20% vs. 85±8%, **Fig. 4F, Supp. Fig. 3B)**. We conclude that simple gating could be sufficient for some cell types, but for others the network needs to be trained on representative data that reflects the issues in the real data. This information can be useful to prioritize labeling efforts to more difficult cells.

Finally, we evaluated the contributions of augmentations by training CellSighter with and without augmentations. We found that removing augmentations reduces the prediction accuracy by 1% to 10%, depending on cell type **(Fig. 4G)**. Expectedly, the effect of augmentations was more significant for cell types that were under-represented in the training set and are difficult to classify, such as stromal cells and dendritic cells. Overall, having a wide plethora of augmentations diversifies the training set and helps overcome imbalances in the prevalence of different cell types.

### CellSighter generalizes across datasets and platforms

We evaluated whether CellSighter could apply to different datasets and platforms. First, we evaluated CellSighter on another MIBI-TOF melanoma dataset that includes 164 0.5×0.5mm^2^ images, altogether comprising roughly 220,000 cells. We trained CellSighter on 132 images and evaluated on 32, where it achieved excellent performance (92±6% **Fig. 5A-B**). We note that this performance is higher than on the lymph node dataset, presumably because cells in this dataset are less dense and the labels underwent extensive manual curation. Next, we turned to a published dataset of Melanoma metastases acquired by Imaging Mass Cytometry (IMC) ^39^. We used the cell classifications provided by the authors to train CellSighter on 55 images and tested the results on 16 heldout images. CellSighter achieved high levels of accuracy (84±17%) on all classes except for stroma (36%), which was mostly confused with tumor cells **(Fig. 5C)**. Since both MIBI and IMC are mass-based multiplexed imaging technologies, we also evaluated the approach on a technology that employs cyclic fluorescence. To this end, we analyzed a published dataset of colorectal cancer acquired using CO-Detection by indexing (CODEX) ^24^. We used the cell classifications provided by the authors to train CellSighter on 112 images (218,372 and tested the results on 28 heldout images **(Fig. 5D)**. Also on this dataset, CellSighter achieved good levels of accuracy (70±19%), albeit lower than the other datasets primarily owing to low performance on several classes.

**Figure 5:**
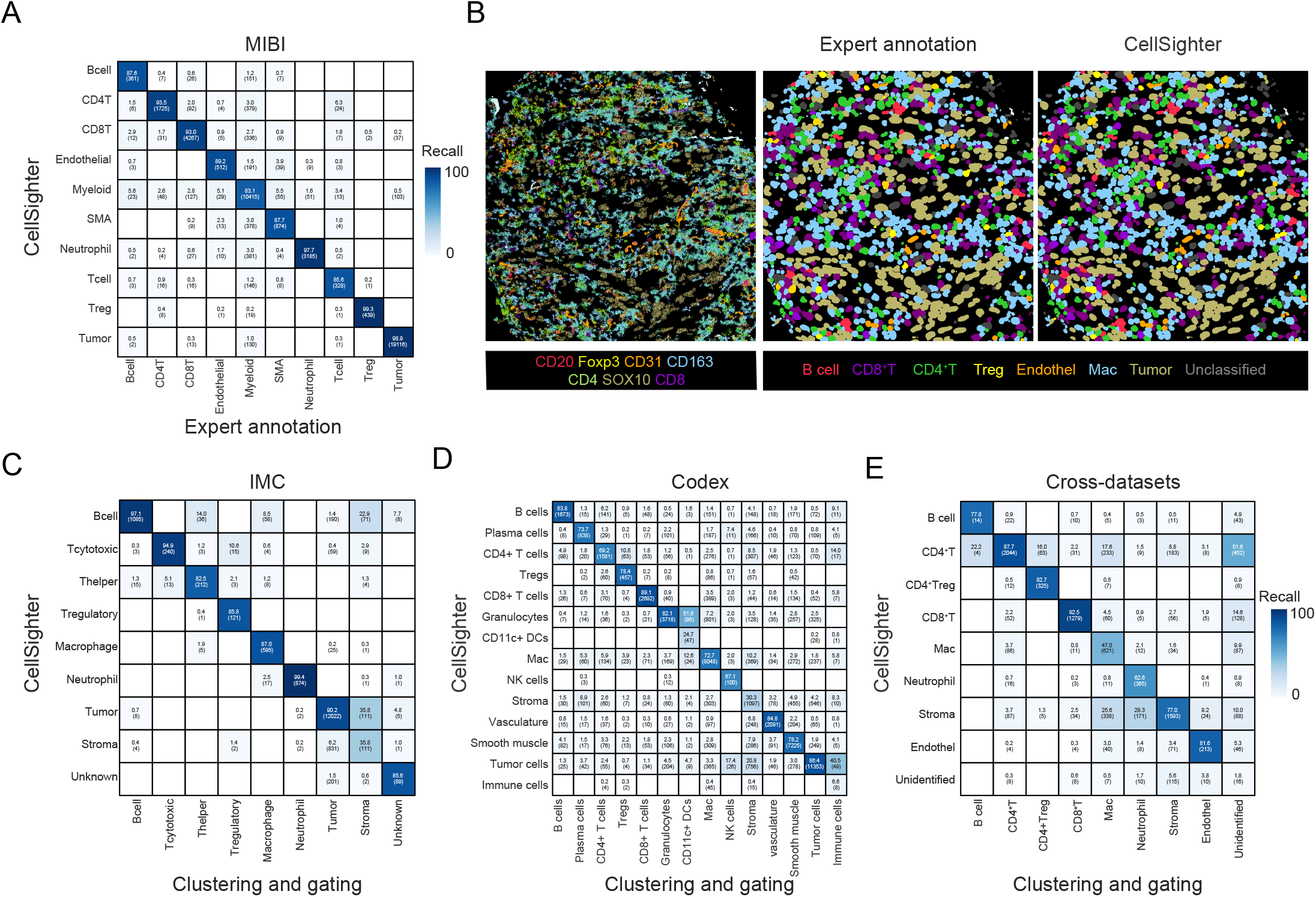
CellSighter generalizes across datasets and platforms. **(A)** Comparison between labels generated by experts (x-axis) and labels generated by CellSighter (y-axis) for a melanoma MIBI dataset. **(B)** For one FOV in the MIBI melanoma dataset, shown are the protein expression levels (left), expert-generated labels (middle) and CellSighter labels (right). **(C)** Same as (A) for IMC data from Hoch et al. ^39^ **(D)** Same as (A) for CODEX data from Schurch et al. ^24^ **(E)** Same as (A) showing performance of the model trained on the lymph node metastases dataset and evaluated on a dataset of the gastrointestinal tract.

Finally, we checked whether a model trained on one dataset could be applied to a different dataset. This is a challenging task since different datasets are collected on different tissues where different populations of cells reside, and these cells may have altered morphology, phenotypes and spatial organizations. Technically, different datasets will typically differ in the number and identity of proteins visualized and may have batch effects relating to the instrumentation and antibodies used at different times. We evaluated how CellSighter, trained on the melanoma lymph node dataset **(Fig.1D)** performed on a dataset profiling the gastrointestinal tract, which shares 19 proteins used to define eight shared cell types **(Fig. 5E)**. We found that training CellSighter on the melanoma lymph node data and evaluating on the gastrointestinal data achieves high results for major cell types that shared all of their defining proteins, including CD4 T cells (88%), Tregs (83%), CD8 T cells (93%) and Endothelial cells (82%). For macrophages the performance was significantly low (47%). Notably, for this class there were unique proteins which were used for expert labeling in one dataset, but not the other. As such, poorer performance could stem from not having enough information for classification, or indicate biological variability in the expression patterns and morphology of these cell types between the lymph node and the gut ^40^.

Altogether, we conclude that CellSighter achieves accurate cell type classification within a single dataset for different types of multiplexed imaging modalities. Across datasets, accurate classification is dependent on having shared lineage-defining proteins, morphology and phenotypes.

## Discussion

Cell classification lies at the heart of analysis of multiplexed imaging, but has hitherto remained labor-intensive and subjective. In this work we described CellSighter, a CNN to perform cell classification directly on multiplexed images. We demonstrated that the network learns both features of protein co-expression as well as spatial expression characteristics, and utilizes both to drive classification. We showed that CellSighter achieves high accuracy (89-90%), on par with current labeling approaches and inter-observer concordance, while drastically reducing hands-on expert labeling time.

CellSighter has several features that we found appealing as users who frequently perform cell classification on multiplexed images. First, CellSighter outputs for each cell not only its classification, but also a confidence score. This type of feedback is nonexistent using current clustering or gating approaches, which often result in variable and arbitrary quality of cell classifications. Working with CellSighter, we found that the confidence scores that are generated are useful in evaluating any downstream analyses that are based on these classifications. Furthermore, evaluating CellSighter’s predictions on a test set is highly informative of label qualities. We consistently found that improving the labeling that is used for training improves CellSighter’s performance and ability to generalize. Therefore, classes that have low prediction accuracies can help the user to identify cell types that are poorly defined. This in turn can direct further efforts to split classes, merge classes, gate or perform manual annotations on the cells of the training set until adequate results are achieved. This process will likely increase the overall labeling quality in multiplexed imaging.

CellSighter also has some caveats that are important to be aware of. Primarily, it is a supervised approach. As such, if a rare cell population (eg Tregs) is not represented in the training set, CellSighter will not be able to identify it in the rest of the dataset. One way to diminish this issue is to validate that for each antibody there are images containing positive staining in the training set. Still, this will not resolve rare populations that are defined based on differential combinations of proteins that are usually associated with more abundant populations. For example, very rare FoxP3^+^ CD8^+^ T cells ^41^ may not be included in the training set and therefore missed. Incidentally, such extremely rare populations are also commonly not classified using standard clustering and gating-based approaches ^24,35,39^. With CellSighter, such cells can be more readily identified by examining cells that were confused between classes. For example, FoxP3^+^ CD8^+^ T cells could be identified by evaluating cells that have high probabilities to be classified as either Tregs or CD8 T cells. The user can then decide whether to add this class and make sure that it is represented in the training set or, alternatively perform subsequent gating of this population.

In addition, CellSighter receives as input images of multiple channels. This has a big advantage in that multiple proteins are assessed simultaneously to call out cell types. For example, CD4 T cell classification will be driven by expression of CD3 and CD4, but not CD8 and assessing these proteins for both the classified cell and its immediate surroundings. This mimics what human experts do when they perform manual labeling and adds to the accuracy of classification. On the flip side, different datasets often include different proteins in their panels. A good example for this is myeloid cells, where different studies measure different combinations of CD14, CD16, MHCII, CD163, CD68, CD206, CD11C etc ^14,24,35,39,42–45^. Transferring models between datasets that don’t share all the proteins used in classification is not straightforward, and reduces the accuracy of classification. There are several solutions to this issue. We found that proteins can be interchanged if they share a similar staining pattern. In addition, as technologies mature, antibody panels will likely increase in size and become more standardized, reducing inter-dataset variability in the proteins measured, which will facilitate transferring labels across datasets and platforms. Altogether, we foresee that in the future, using machine-learning approaches such as CellSighter will streamline data integration, such that knowledge would transcend any single experiment and consolidate observations from different studies ^46,47^. While CellSighter is undoubtedly not there yet, it is an important step in facilitating this process.

## Methods

### Datasets and expert annotations

#### MIBI Melanoma lymph node dataset

The dataset contains sixteen 0.8×0.8mm^2^ images, altogether encompassing 116,808 cells. Of these, the cells from 12 images were used for training and the model was evaluated on the remaining four images. Labels for all cells were generated using FlowSOM clustering ^22^, in combination with gating and sequential rounds of visual inspection and manual annotation. The following populations have been defined: B cells (CD20, CD45, CD45RA), DCs (DC_SIGN, CD11c, CD14, CD45, CCR7, CD4), CD4 T cells (CD4, CD3, CD45), T regs (CD4, FoxP3, CD3, CD45), Macrophages (CD45, CD68, CD163, CD206, DC_SIGN, CD14, CCR7), CD8 T cells (CD8, CD3, CD45, Granzyme B), Stroma (COL1A1, SMA), Follicular Germinal B cells (CD20, CD21, CD45, CD45RA,), HEVs (CD31, Meca79), Memory CD4 Tcells (CD4, CD3, CD45, CD45RO), NK cells (CD45, CD56), Neutrophils (S100A9_Calprotectin), endothelial cells (CD31) and Tumor (MelanA, SOX10). The following expert-annotated classes were not included in predictions: unidentified, which contains a mixture of various proteins, and CD3-only, together encompassing 5% of the data. Expression of all proteins across cell types can be found in supplementary figure 3C.

#### MIBI Melanoma dataset

174 0.5×0.5mm^2^ images, altogether encompassing 220,016 cells. Labels for all cells were generated using sequential gating and sequential rounds of visual inspection and extensive manual annotation. Of these, the cells from 132 images were used for training and the model was evaluated on the remaining 42 images. The following populations have been defined: B cells (CD20, CD45), CD4 T cells (CD3, CD4, CD45), CD8 T cells (CD3, CD8, Granzyme B, CD45), Endothelial (CD31), Myeloid (CD16, CD68, CD163, CD206, DC_SIGN, CD11c, CD45,), SMA (SMA), Neutrophil (MPO_ Calprotectin), T cell (CD3 only), T regs (FoxP3) and Tumor (SOX10). Expression of all proteins across cell types can be found in supplementary figure 3D.

#### MIBI GastroIntestinal (GI) dataset

Labels for eighteen 0.4×0.4mm^2^ images were generated using FlowSOM clustering ^22^, in combination with pixel clustering ^31^ and sequential rounds of visual inspection and manual annotation. The model trained on the melanoma lymph node dataset was ran on 9,532 cells from the following classes, which were shared across the two datasets: T regs (FoxP3, CD4, CD3, CD45), CD8 T cells (CD8, CD3, GranzymeB, CD45), CD4 T cells (CD4, CD3, CD45), B cells (CD20, CD45RA, CD45), Macrophages (CD68, CD206, CD163, CD14, DC-SIGN, CD45), Neutrophil (S100A9-Calprotectin), Stroma (SMA, COL1A1) and Endothelial (CD31). Expression of all proteins across cell types can be found in supplementary figure 3E.

#### IMC Melanoma dataset

Data and cell classifications were taken from Hoch et al. ^39^. Of these, the cells from 55 images were used for training and the model was evaluated on the remaining 16 images, altogether encompassing 70,439 labeled cells. The following populations, as defined by the authors, have been used: Tumor, B cell, CD4 T cells, Macrophage+pDC, CD8 T cells, Stroma, Neutrophil, Tregs and unknown. Annotations were provided for a subset of cells. For this dataset, the crop size for the CNN was chosen to be 30×30 pixels because of the image resolution. To train CellSighter the following subset of protein channels was used: CD4, CD20, SMA, SOX10, FOXP3, CD45RO, Collagen I, CD11c, CD45RA, CD3, CD8a, CD68, CD206/MMR, S100, CD15, MPO, HLA-DR, CD45, CD303, Sox9, MiTF, CD19, p75

#### CODEX Colorectal dataset

Data and cell classifications were taken from Schurch et al. ^24^. Of these, the cells from 112 images were used for training and the model was evaluated on the remaining 28 images, altogether encompassing 218,372 cells from the following classes: Macrophages (CD68+CD163+ macrophages, granulocytes, CD11b+CD68+ macrophages, CD68+ macrophages, CD163+ macrophages, CD68+ macrophages GzmB+), DCs (CD11c+ DCs), CD4 T cells (combined with CD4+ T cells GATA3+ and CD4+ T cells CD45RO+), CD8 T cells, T regs cells, Granulocytes, Vasculature, Stroma, Smooth muscle, Plasma cells, B cells, Tumor cells, Immune cells, NK cells. We chose to not work on the other classes in the dataset due to the fact that they were either not well defined or that they don’t have a lot of samples. To train CellSighter the following subset of protein channels was used: CD7, GATA3, CD44, FOXP3, CD8, p53, CD45, beta-catenin, HLA-DR - MHC-II, CD45RA, CD4, CD21, MUC-1, CD20, Na-K-ATPase, Cytokeratin, CD11b, CD56, aSMA, CD11c, Granzyme B, CD15, Synaptophysin, GFAP, CD3, Chromogranin A, CD163, CD45RO, CD68, CD31, Podoplanin, CD34, CD38, CD138.

### CellSighter

CellSighter is an ensemble of CNN models, each based on a ResNet50 backbone ^48^. The input for each model is a 3-dimensional tensor, consisting of cropped images of K proteins centered on the cell to be classified, a binary mask for the cell to classify (1’s inside the cell and 0’s outside the cell), and a similar binary mask for all the other cells in the environment **(Fig. 1C)**. The crop size can vary, but ideally should include the cell and its immediate neighbors. For the datasets at hand no significant differences were observed when varying the crop sizes from 40 – 100 pixels (corresponding to ±20-50 μm^2^, **Supp. Fig. 1B**). A crop size of 60×60 pixels was used for all datasets except the IMC, where a crop size of 30×30 pixels was used to account for the different resolution. To account for class imbalance, rare classes were upsampled such that the major lineages (Myeloid, T cell, tumor, B cell, other) were represented in equal proportions. *Training*: In training, the images randomly undergo a subset of the following augmentations: no augmentation, rotations, flips, translations of the segmentation mask, resizing of the segmentation mask by up to 5 pixels, shifts of individual protein channels in the X-Y directions by up to 5 pixels with probability of 30%, and gaussian signal averaging in a window of 5 pixels followed by Poisson sampling. The final cell classification is given by integrating the results from ten separately-trained CNNs. We found that randomization in initializations and augmentations are sufficient to generate sufficient diversity between the models. Each model predicts the probability of each class for the cell. Those probabilities are then averaged to generate one probability per cell type. The final prediction is the class with the maximal probability and the prediction confidence is the probability value.

The code for CellSighter can be found at: https://github.com/KerenLab/CellSighter

### XGBoost

Python’s XGBoost gradient boosting tree model was used for benchmarking experiments on tabular data (https://github.com/dmlc/xgboost ^38^). Input to the model included the arcsinh transformed expression values per cell normalized by cell size for all the proteins that were used to train CellSighter ^18^. The model was trained with the following parameters: n_estimators=100 and max_depth=2, the rest of the parameters were kept as the default of the library.

### Gradient analysis

The analysis was performed using one of the models of the ensemble on 5000 cells from the test set. Small cells (<50 pixels) were removed since their size is too small for spatial analysis, leaving 4961 cells. For each of these cells, the sum of positive gradients for each marker was calculated in concentric circles centered in the middle of the input, ranging from 1 to 18 pixels in jumps of 2. For each cell the radius of the circles is then normalized by the radius of the cell. Gradients are normalized relative to the largest radius profiled.

### Neutrophil analysis

#### Calprotectin patch analysis

Calprotectin patches were obtained by dilating the signal using a kernel of size 3 and identifying connected components. Cells that overlap with Calprotectin patches are identified.

#### Simulations

All 237 Neutrophil cells from the test set were used. Calprotectin signal was simulated using Poisson sampling with lambda=4 normally distributed with a standard deviation of 5 pixels. Simulations were performed in which this signal was moved relative to the center of the cell ranging from 0 to 15 pixels in x and y directions.

### CD4/CD20 simulations

#### CD20 patch experiments

549 CD4 T cells were randomly-sampled and their cognate CD4 and CD20 signals removed to avoid any confounding factors incurred by the original signals. CD4 signal was simulated by Poisson sampling with lambda=1.3 in horizonal and vertical distances of at most 5 pixels from the border of the cell segmentation. CD20 was simulated around a random point on the border of the cell with Poisson sampling with lambda=1.3 with a uniform distance between 0 and 6. Cells that are smaller than 15 pixels across their minor axis were filtered to eliminate complete overlap of the patch with the cell.

#### CD20 membrane experiments

200 CD4 T cells were randomly-sampled and their cognate CD4 and CD20 signals removed to avoid any confounding factors incurred by the original signals. CD4 signal was simulated by Poisson sampling with lambda=1.3 in horizonal and vertical distances of at most 5 pixels from the border of the cell segmentation. CD20 was simulated similarly, but varying the percent of the membrane that is covered by the signal to be 12.5%, 25%, 50% and 100%. In order to keep the overall signal intensity in the cell similar, the number of sampled points increased proportionally to the decrease in membrane size.

### Training with different input sizes

For these experiments, 100, 250, 500, 1000 or 5000 cells were randomly sampled from the training set for each class. In cases in which there were not enough cells in the data for sampling (e.g. Tregs had only 440 cells in the training set), all cells from that class were sampled, but results for these values for these classes are not reported to allow comparisons across experiments. Figures show the mean and std of five independent experiments.

### Landmark cell analysis

Landmark cells were defined for each class as the cells that express above 20^th^ percentile of the proteins that define the class, and are not above the 15^th^ percentile value of expression of any other protein. E.g. landmark tumor cells strongly expressed either SOX10 or MelanA and no other lineage protein. For the following classes some deviations from this formulation were necessary to allow enough cells for training: For myeloid cells the threshold for the other markers was the 20^th^ and not 15^th^ percentile. For Tregs and neutrophils only high expression of FoxP3 and Calprotectin was used respectively, without consideration for other proteins. Visual inspection validated that these cells were indeed landmarks of their classes.

Landmark cells were partitioned to train and test by the same image partition as in figure 1. CellSighter was trained on the landmark training cells and tested both on the landmark test cells **(Supp. Fig. 3A)** and on all cells in the test **(Supp. Fig. 3B)**.

## Acknowledgements

The authors thank Tal Keidar Haran, Eli Pikarsky and Michal Lotem for samples and data of melanoma lymph metastases. L.K. is supported by the Enoch and Azrieli foundations, and grants funded by the Schwartz/Reisman Collaborative Science Program, European Research Council (94811), the Israel Science Foundation (2481/20, 3830/21) within the Israel Precision Medicine Partnership program. I.M is supported by a EU - Horizon 2020 - MSCA Individual Fellowship (890733). S.B. is a Robin Chemers Neustein AI Fellow and acknowledges funds from the Carolito Stiftung and the NVIDIA Applied Research Accelerator Program.

## Notes

https://github.com/KerenLab/CellSighter

### Competing Interest Statement

The authors have declared no competing interest.

