## Supplemental figures for "CellSighter – A neural network to classify cells in highly multiplexed images"

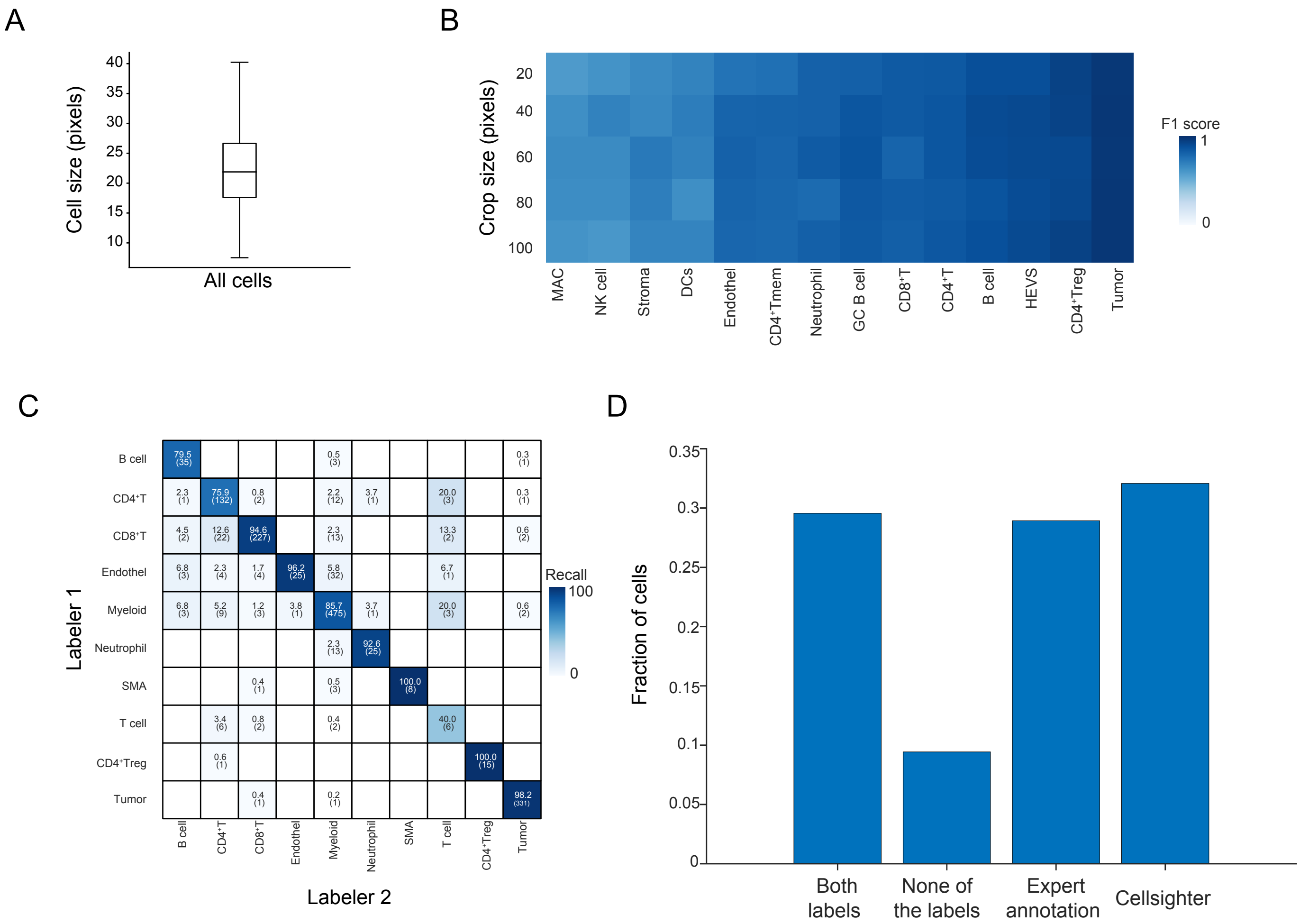


**Supplementary Figure 1**

**(A)** Boxplot showing the major axis length in pixels for all cells in the Melanoma lymph node metastases dataset. **(B)** Shown is the agreement between CellSighter and expert labeling (F1 score, blue) for different cell classes (x-axis) when varying the crop size of the input into CellSighter between 20 and 100 pixels. **(C)** For the melanoma dataset, shown is the agreement on the labels of all cells from a single FOV between two different human labelers. **(D)** Expert inspection of cells differing in classification between CellSighter and expert annotation (cells off the diagonal in figure 5A). Inspection was performed blindly, without knowledge of the source of the label.

Figure S2

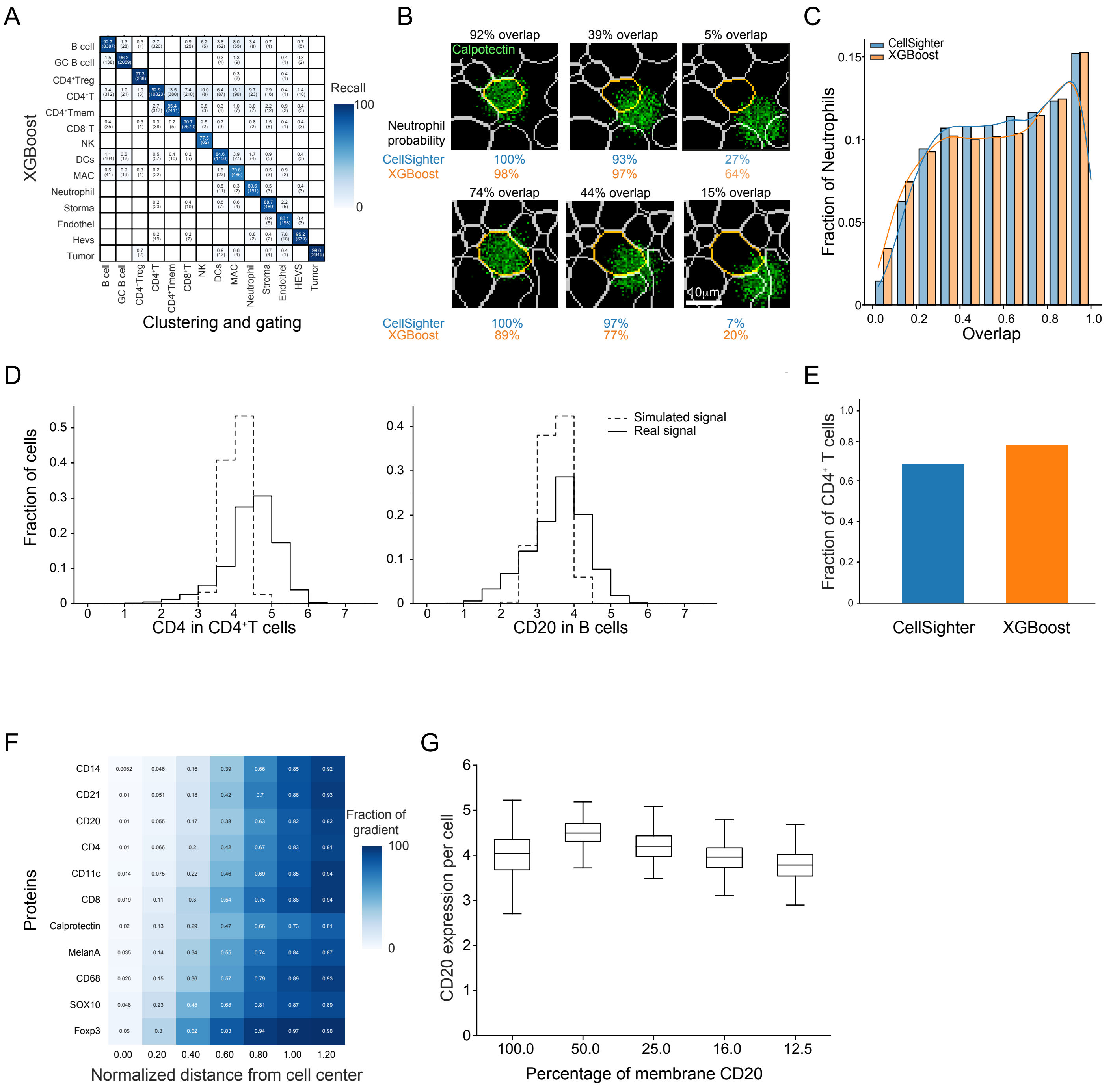


**Supplementary Figure 2**

**(A)** Comparison between labels generated by experts (x-axis) and labels generated by running gradient-boosting on the expression matrix (XGBoost, y-axis). **(B)** Example images from simulations in which the level of intersection of Calprotectin (green) with a cell-to-be-classified (yellow) is varied. Shown are three degrees of overlap for two cells. **(C)** Results of simulations for 237 cells with different overlaps of Calprotectin as shown in (B). Shown are the proportion of cells classified as Neutrophils (y-axis) at different levels of intersection of their Calprotectin signal with the cell segmentation (x-axis). At low intersections, CellSighter (blue) classifies less cells as Neutrophils than XGBoost (orange). **(D)** Histogram showing expression levels of CD4 in CD4 T cells (left) or CD20 in B cells (right) for real (black) and simulated (gray) data. **(E)** Crops centered on CD4 T cells had their cognate CD4 and CD20 signals removed and CD4 was reintroduced as a membranous signal. Shown are results for the fraction of these cells that were classified as CD4 T cells (y-axis) by CellSighter and XGBoost (x-axis). **(F)** Shown is the normalized sum of gradients (y-axis) as a function of the normalized integrated distance from the cell center (x-axis) for different lineage proteins. The gradients for each protein were evaluated in its respective cell type: CD14 – Macrophages, CD21 – Follicular germinal B cells, CD20 – B cells, CD4 – T helper cells, CD11c – DCs, CD8 – CD8 T cells, Calprotectin – Neutrophils, MelanA – Tumor cells, CD68 – Macrophages, SOX10 – Tumor cells, Foxp3 – Tregs. **(G)** For cells in which CD20 was simulated as a membranous signal with varying overlap with the membrane, shown is the CD20 expression per cell (y-axis) as a function of the percentage of membrane CD20 (x-axis).

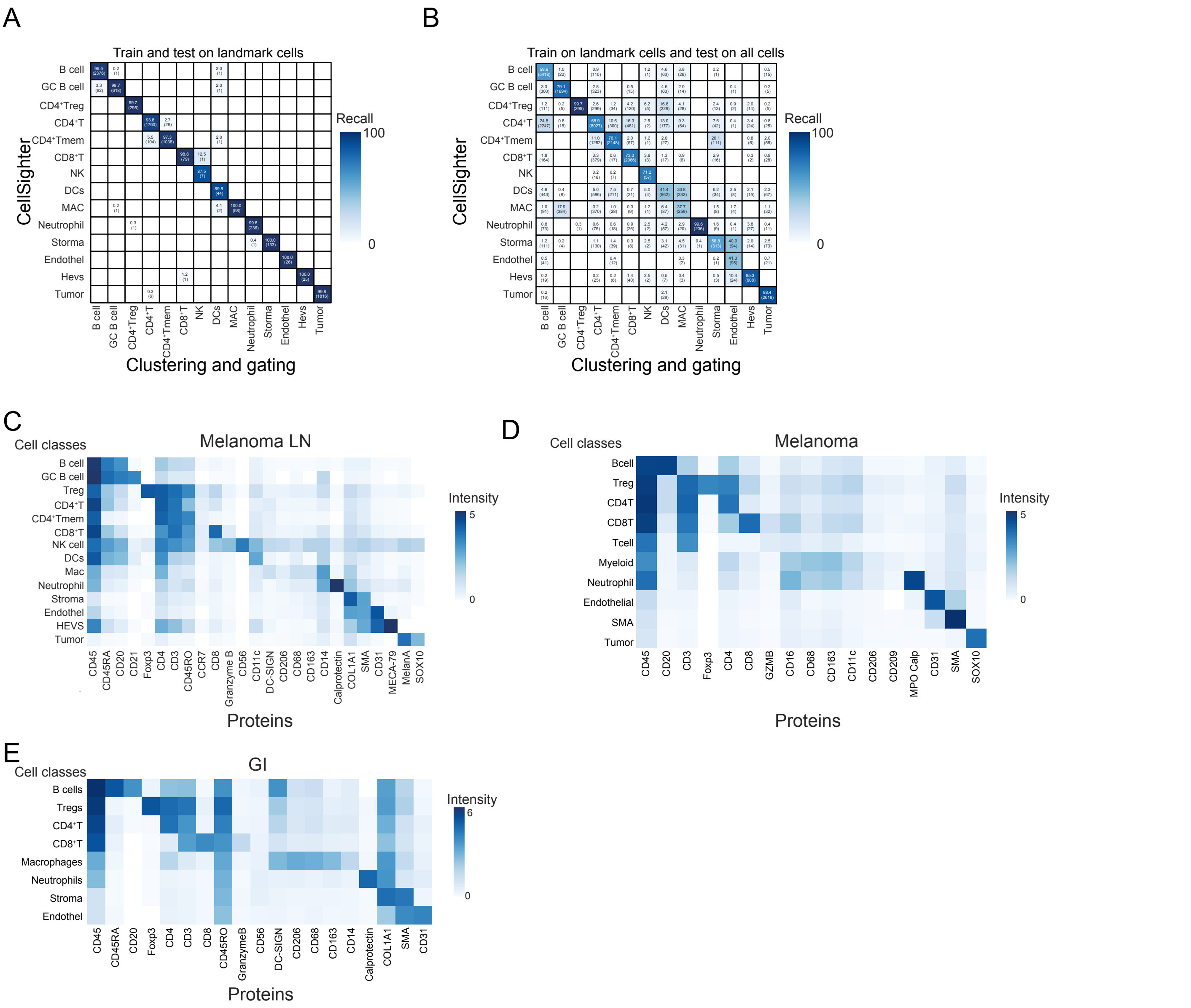


**Supplementary Figure 3**

**(A)** Comparison between labels generated by experts (x-axis) and labels generated by running CellSighter (y-axis). Shown are results for a model that was trained and evaluated on landmark cells. **(B)** Comparison between labels generated by experts (x-axis) and labels generated by running CellSighter (y-axis). Shown are results for a model that was trained on landmark cells and evaluated on the complete dataset. **(C)** For the melanoma lymph node data, shown is expression of distinct proteins (x-axis) across cell classes (y-axis). **(D)** For the melanoma data, shown is expression of distinct proteins (x-axis) across cell classes (y-axis). **(E)** For the gastrointestinal data, shown is expression of distinct proteins (x-axis) across cell classes (y-axis).
